# Attentional reconfiguration during acute and sustained fear

**DOI:** 10.1101/2025.11.06.686932

**Authors:** Kerttu Seppälä, Nella Rantala, Matthew Hudson, Vesa Putkinen, Severi Santavirta, Jukka Hyönä, Lauri Nummenmaa

## Abstract

**BACKGROUND:** Fear is a fundamental survival mechanism that both enhances the chances of survival by rapid detection and adaptive responses to potential threat and by optimizing sensory input and cognitive processing. Here we used naturalistic design with eye tracking to map the spatiotemporal dynamics of attention and arousal during fearful events with slow and fast temporal dynamics.

**METHODS:** 21 participants watched a full-length horror movie while their eye-movements were recoded with eye tracker. Moment-to-moment intensity of sustained fear as well as the onsets of the jump scares (acute fear) were annotated and used to predict gaze parameters (fixation duration & counts, blink frequency, saccade amplitude, pupil size and intersubject synchronization of gaze position).

**RESULTS:** Acute fear events led to shortening of fixation duration, suppression of blinking, as well as increase of fixation count, saccade amplitude, and pupil size. Sustained fear was in turn associated with increased pupil size and decrease in blinking and saccade amplitude. These effects remained significant when controlling for luminosity.

**CONCLUSIONS:** During natural vision both acute and sustained fear cause rapid reconfiguration, state dependent and individual changes in visual attention prioritizing that is accompanied by increased affective arousal.

## Introduction

Fear acts as survival intelligence, constantly scanning and responding to potential threats, and managing their efficient processing in the sensory and cognitive systems to maximize survival odds. Numerous studies have established that attentional processing of threats is prioritized (Vuilleumier 2005), which is reflected in eye-tracking data as selective attentional orienting to and engagement with the threat signals (Nummenmaa, Hyona et al. 2006, Nummenmaa, Hyönä et al. 2009). However, mapping the attentional dynamics of fear is difficult due to the complex spatiotemporal dynamics of eye movement guidance. Establishing the links between eye movements and the contents of a high-dimensional stimulus is challenging, and prior research has often relied on the analysis of spatial gaze data in response to artificial static snapshots of scenes in pictures. This however precludes quantifying the effects of temporally varying stimulus properties, such as moment-to-moment visual saliency or emotional valence, on gaze control. It has thus been debated whether the results from such simplified conditions transfer to natural vision operating in the dynamic social environment (Williams and Castelhano 2019), particularly as the visual system is differently influenced by static versus dynamic stimuli (Dorr, Martinetz et al. 2010). Furthermore, the fear system operates at multiple time scales, from trying to reduce uncertainties and preparing to respond to distal threats to actual fight-or-flight responses during proximal dangers (Mobbs, Headley et al. 2020). Although the neurobiology behind the sustained and acute threat systems have been mapped extensively in prior studies (Mobbs, Petrovic et al. 2007, Mobbs, Yu et al. 2010, Hudson, Seppälä et al. 2020), the effects of acute and sustained fear on gaze behaviour during dynamically evolving, naturalistic fear remain poorly characterized.

Horror movies are excellent means for evoking strong, temporally varying fear in humans (Nummenmaa 2024). Laboratory studies have confirmed that viewing cinema is a reliable means for inducing strong and consistent self-reported fear (Gross and Levenson 1995, Schaefer, Nils et al. 2010). Compared with pictures or sounds, threatening audiovisual stimuli also cause stronger autonomic activation, as indicated by increased heart rate (Schaefer, Nils et al. 2010, Bhushan and Asai 2018). Eye tracking studies using cinematic fearful stimuli show increased blinking rate (Bhushan and Asai 2018, Maffei and Angrilli 2019) thereby reducing sensory input (Maffei and Angrilli 2019) since with every blink visual information is lost (Nakano, Yamamoto et al. 2009) maybe leaving more time for survival solution. Gaze position, on the other hand, synchronizes across individuals during cinema viewing (Henderson 2003, Lahnakoski, Glerean et al. 2014, Nummenmaa, Smirnov et al. 2014), and particularly during strongly emotional scenes (Subramanian, Shankar et al. 2014), suggesting that emotions may provide a bottom-up mechanism for gaze guidance to promote swift response to threats. Further, studies on saccade parameters are often limited to the anxious or depressed patients who show different patterns as healthy volunteers (Nakano, Yamamoto et al. 2009, Chen, Clarke et al. 2014, Gao, Xin et al. 2023). Pupil size in turn fluctuates constantly even in rest (Wilhelm, Lüdtke et al. 1998), but adapts primarily to adjust the amount of light landing on the retina (Sirois and Brisson 2014). However, emotional arousal (Lowenstein, Feinberg et al. 1963, Bradley, Miccoli et al. 2008, Van Steenbergen, Band et al. 2011), fear (Leuchs, Schneider et al. 2017), and even imagined emotional arousal (Henderson, Bradley et al. 2018) cause the pupil to dilate. These luminance-independent pupillary responses are mediated by the adrenergic and cholinergic systems engaged during emotions to maximize attentional prioritization and arousal (Joshi, Li et al. 2016, Reimer, McGinley et al. 2016).

In addition to investigating the effects of slow, phasic threat, horror movies allow evoking abrupt fear through jump scares where the threat emerges suddenly and unpredictably. Most prior work on such acute fear responses is based on the fear conditioning paradigm e.g. (Korn, Staib et al. 2017). In fear conditioning paradigms, pupil reacts to arousing inputs by dilating in an event-related fashion over the time course of a few seconds (Kluge, Bauer et al. 2011, Wiemer, Mühlberger et al. 2014, Korn, Staib et al. 2017). Fixations in turn become longer and fewer in numbers following threat versus safety signals (Klein, Ginat-Frolich et al. 2021, Rodriguez-Sobstel, Wake et al. 2023), while saccade duration remains unchanged (Deuter, Schilling et al. 2013), even in real-life scenarios such as a fear of falling (Naranjo, Cleworth et al. 2017). Altogether, these data show the feasibility of measuring eye parameters following acute threats. However, in comparison with conditioned stimuli, horror movies with jump scares would allow investigation of the consequences of natural, unconditioned threats on the oculomotor system.

Against this background, horror movies serve an excellent testbed for measuring the effects of fear on arousal and attentional guidance in naturalistic settings. First, modelling the associations between temporal unfolding of fear and different gaze parameters (e.g. fixation and blink rates, pupil size) enables the study of slow-frequency oculomotor coupling between fear and different aspects of gaze guidance under natural dynamic vision. Here we show how slow-frequency fluctuations in sustained fear are consistently linked with pupil dilation, shortening of saccades and reduced number in blinks, indicative of increased arousal and vigilance. Second, brief eye movement responses evoked by the rapid-onset “jump scares” allow for modelling of the high-frequency coupling between acute unconditioned fear and gaze control. Acute threats in turn led to rapid pupil dilation, shortening of fixations and reducing the number of blinks, yet to increased number of fixations and larger saccade amplitudes. Gaze synchronization between the subjects, however, did not change, indicating that at the point of abrupt fear, attention is drawn to scanning and responding to potential threats.

## Materials and methods

### Subjects

Twenty-eight participants (21 women, mean age 27.6 (SD 8.4), BMI 23.7 (SD 0.94) completed the study. Four had corrected vision. Participants were recruited through social media and email lists, and they completed informed consent forms prior to participating. They were compensated with movie tickets or study credits. All participants successfully finished the eye-tracking study. Two participants did not complete all personality questionnaires and were therefore excluded from the personality trait correlation analysis (see SI file). An additional 20 participants, who did not take part in the main experiment, provided dynamic ratings of fear intensity while watching the movie, and the averaged moment-to-moment fear intensity was used as a predictor of the eye-tracking measures (see below).

### Movie stimulus and dynamic fear ratings

The stimulus was the movie Conjuring 2, directed by James Wan (Wan 2016). This movie was chosen because it is one of the highest-rated horror movies in online databases (Rotten Tomatoes, International Movie Database, AllMovie) and our pilot survey with 7 participants confirmed that the movie was in general evaluated scary, as it contained frequent jump scares (acute threat onsets) allowing modelling of both sustained and acute fear responses (Hudson, Seppälä et al. 2020). The movie was shortened by 24 min 42 s resulting in 1 h 49 min 24 s for watching, and was then split into 30 segments with Apple iMovie. Each segment lasted 2-4 min. Cut points followed the existing cuts and the natural flow of the plot. In a separate experiment, 20 subjects not participating in the eye tracking study evaluated their moment-to-moment experience of intensity of fear during the movie by moving a slider on a visual analogue scale (Hudson, Seppälä et al. 2020), as shown in Figure 1 with sample images of the movie. To assess the effects of acute threats on the visual system, we extracted the timings of the 22 jump scares in the movie from online database Where is the Jump? (http://wheresthejump.com, 2017) and constructed a stick function regressor for their onsets.

**Figure 1.**
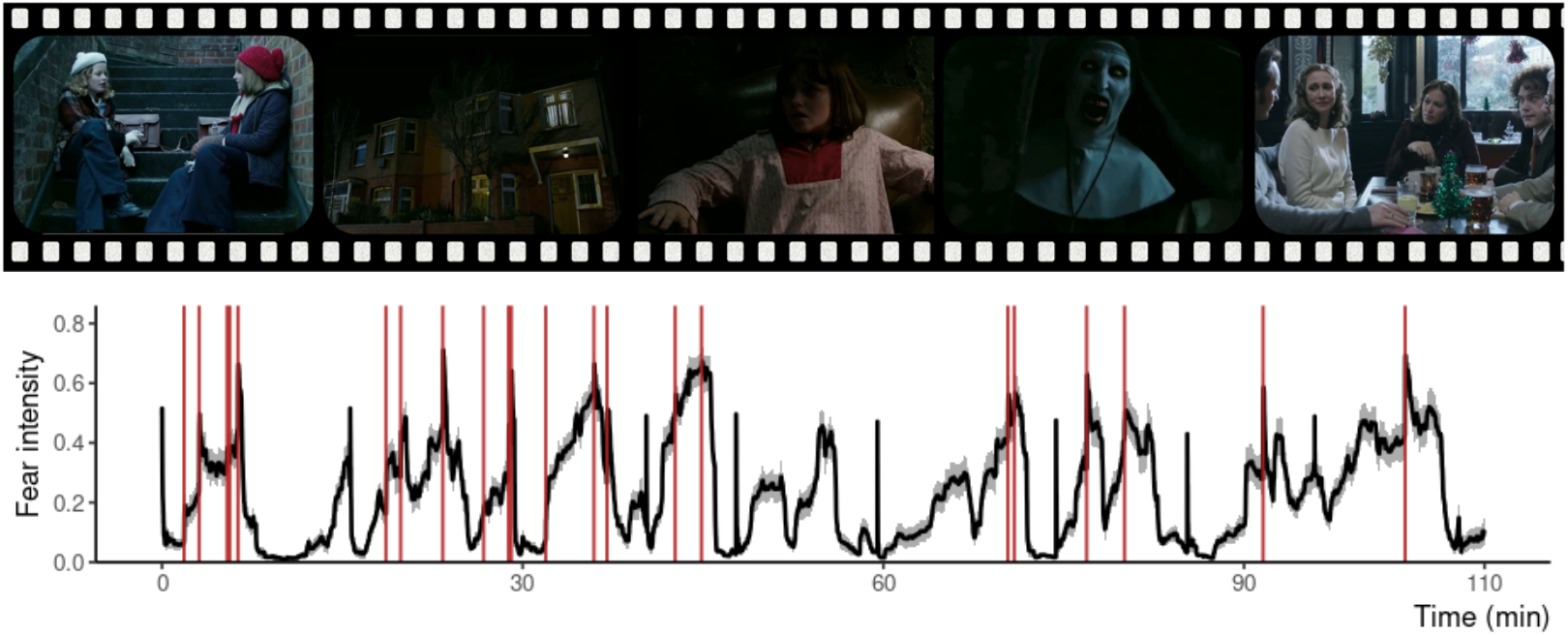
Top: Sample frames from the movie with neutral and scary contents. Bottom: Mean and standard error of the mean (SEM) for fear ratings of 20 subjects, 1 indicates maximal fear, 0 no fear at all. The red vertical lines mark the jumps scares (abrupt threats).

### Eye movement recordings

Eye movements were recorded with EyeLink 1000+ tracker (SR Research Ltd., Mississauga, Ontario, Canada) with 500 Hz temporal resolution using 16 mm lens. The 9-point calibration was accepted if the average error was below 1°. Illuminator power was set at 75 %, and the lightning of the experiment room was kept constant across the subjects. The stimulus was presented on a 26’’ screen (16:10) with a resolution of 1000×720 pixels, one segmented part of the movie at a time. Participants were seated at 65 cm away from the eye tracker. Drift correction was performed before each trial (i.e. each of the 30 movie segments), and the eye tracker was calibrated at the beginning of the experiment and after every third segment with 9-point calibration. Participants were instructed to watch the movie as they would be watching TV at home, but were encouraged to avoid movements during the experiment.

### Self-reports

After every three movie segments, the participants reported their feelings of happiness, anger, disgust, sadness, surprise, anxiety, and fear on a scale of 0-100. After the experiment, the subjects filled in-house questionnaires about the movie (immersion, emotion regulation during scary scenes and content of the movie, horror movies in general, somatic sensations during the movie (e.g. sweating, shivers, heart racing etc.), and the experience of fear, and personality trait questionnaires (see the SI file).

### Preprocessing of the eye movement data

First, the time series for pupil size, fixation duration and coordinates, saccade amplitudes, and blinks were extracted using the SR Research Data Viewer tool. Data were preprocessed and analysed using Matlab 2020b and RStudio (R Core Team (2023). R: A Language and Environment for Statistical Computing (R Foundation for Statistical Computing, Vienna, Austria. https://www.R-project.org/). To generate a continuous time series for **pupil size**, the pupil size during each fixation was extrapolated until the next fixation. The **fixation rate** time series was constructed by first removing fixations shorter than 80 ms and then time series was converted to Hz (fixations per second). The **saccade amplitude** time series was constructed as step function with amplitude as value for the length of the saccade, leaving intermitting fixations with 0. The **blink** time series was constructed as a binary time series where blink onsets were stored as stick functions and converted to Hz (blinks per second). For all the time series, data points more than 2,5 SD of the mean value were considered as outliers and were replaced with the previous value with Matlab function Filloutliers. **Intersubject correlation (e-ISC)** of gaze position was calculated with e-ISC-toolbox for Matlab (Nummenmaa, Smirnov et al. 2014), where mean spatial correlation was estimated on participant-wise moment-to-moment fixation heatmaps. Finally, the eye-tracking data were smoothed with a moving window of 200 ms for analysis (5 s smoothing for plotting). To control for low-level visual characteristics of the movie, a timeseries for framewise luminosity was computed using root mean square (RMS) of **relative luminance** with spectral weight of human vision. The timeseries for **intensity of fear** was constructed as described previously in (Hudson, Seppälä et al. 2020).

### Data analysis Eye movement responses to sustained fear

We first computed bivariate Pearson correlations between eye movement time series (saccade amplitude, fixation duration, pupil size, e-ISC, and the number of blinks or fixations per second), luminosity and intensity of fear time series. To control for the potential effect of the luminance on gaze parameters, a complementary methodological approach was used. First, a subject-wise least-squares linear regression model was fitted to each eye tracking time series using relative luminance as the predictor. Subsequently, the subject-wise residual time series (containing all variation not explained by luminosity) were predicted by the intensity of fear in separate linear regression models. Subject and model-wise standardized regression coefficients (betas) were extracted, and their distribution was tested against zero using one-sample t-tests to reveal which eye movement parameters were consistently associated with fear at the sample level.

### Eye movement responses to acute fear

To model the evoked responses to jump scares, we first extracted the eye movement time series around each jump scare (from 1 s before to 5 s after the jump scare). The evoked responses were then baseline-corrected to the 1s prior to the jump scare. To estimate whether the gaze parameters deviated significantly from baseline after a jump scare, we applied a cluster-based permutation test using a non-parametric sign-flipping approach, as implemented in cluster-based permutation testing frameworks developed for EEG/MEG studies (Maris and Oostenveld 2007). A one-sample t-test was conducted at each time point across subjects to test whether the mean signal differed from zero. Time points exceeding a critical t-threshold (corresponding to a two-tailed test with α = 0.05) were considered supra-threshold. Spatially contiguous supra-threshold points were grouped into temporal clusters, and for each cluster, the sum of absolute t-values was computed. To build a null distribution of cluster-level statistics under the null hypothesis, 1000 permutations were performed using a random sign-flipping procedure. Supra-threshold time points were clustered, and the maximum cluster statistic was recorded for each permutation. This yielded a null distribution of maximal cluster statistics. The p-value for the observed cluster was computed as the proportion of permutations in which the maximum cluster statistic under the null exceeded the observed statistic. A cluster was considered statistically significant if its statistic exceeded the 95th percentile of the null distribution (p < 0.05, two-tailed). This procedure inherently controls for multiple comparisons.

## Results

### Self-reports

Figure 2 shows the time series for the self-reported emotions while viewing the movie. These data confirmed that the movie successfully elicited the targeted emotions. On average, subjects experienced high levels of fear, anxiety and pleasure, moderate levels of disgust and tranquility, and low levels of anger, confusion, joy, and sadness.

**Figure 2.**
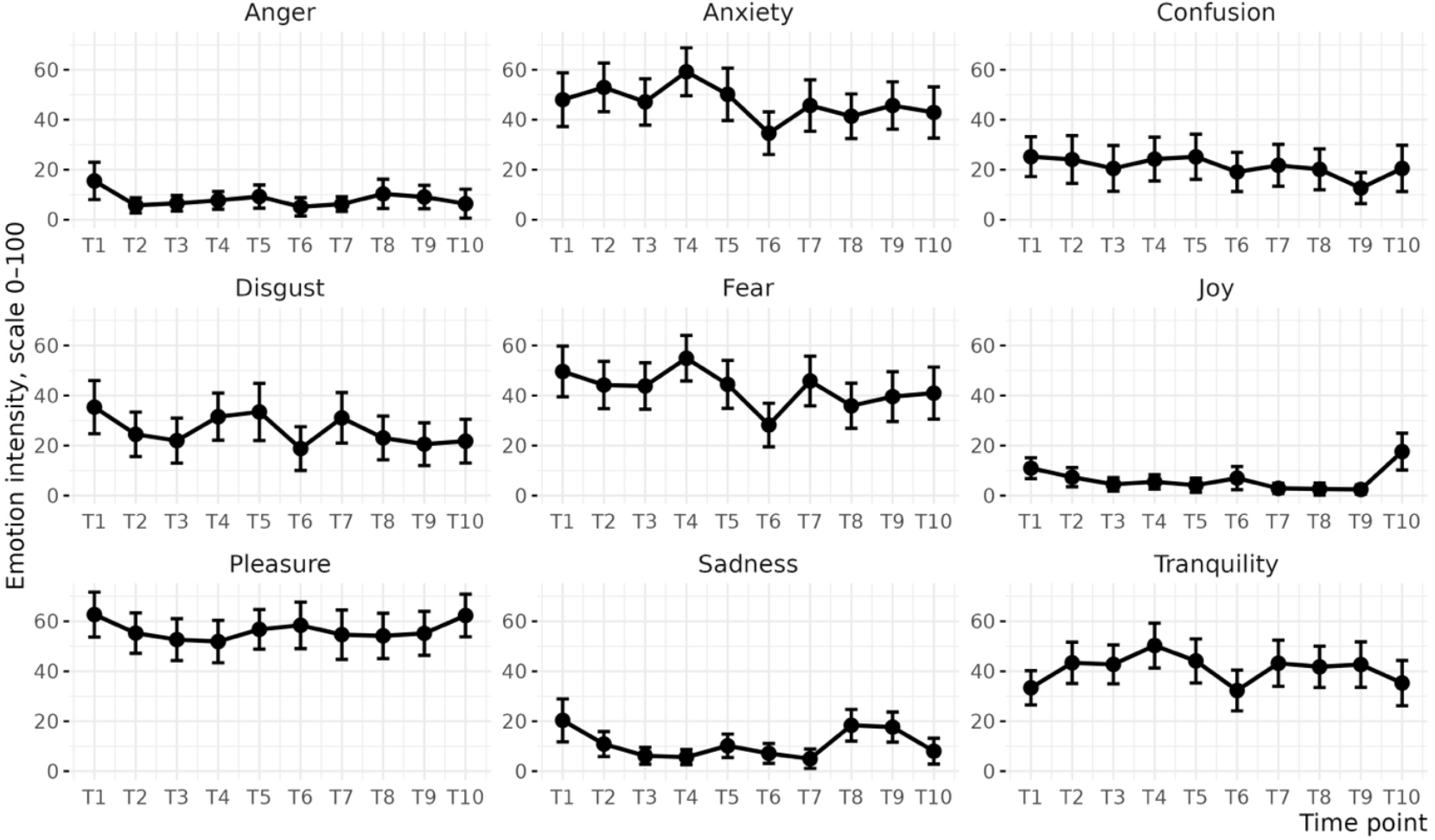
Means and 95% confidence intervals for the experienced emotions on a scale 0-100 (0 = not at all, 100 = the strongest possible feeling) while viewing the movie.

### Effects of sustained fear on oculomotor and pupillary responses

The mean time series (with SEM) and the distributions of intensity of fear, luminance, and eye movement parameters are shown in Figure 3. These indicated that the intensity of fear varied significantly throughout the movie, with concomitant fluctuations in the gaze parameters. Eye movements were also moderately synchronized across subjects throughout the experiment, with mean e-ISC exceeding 0.3.

**Figure 3.**
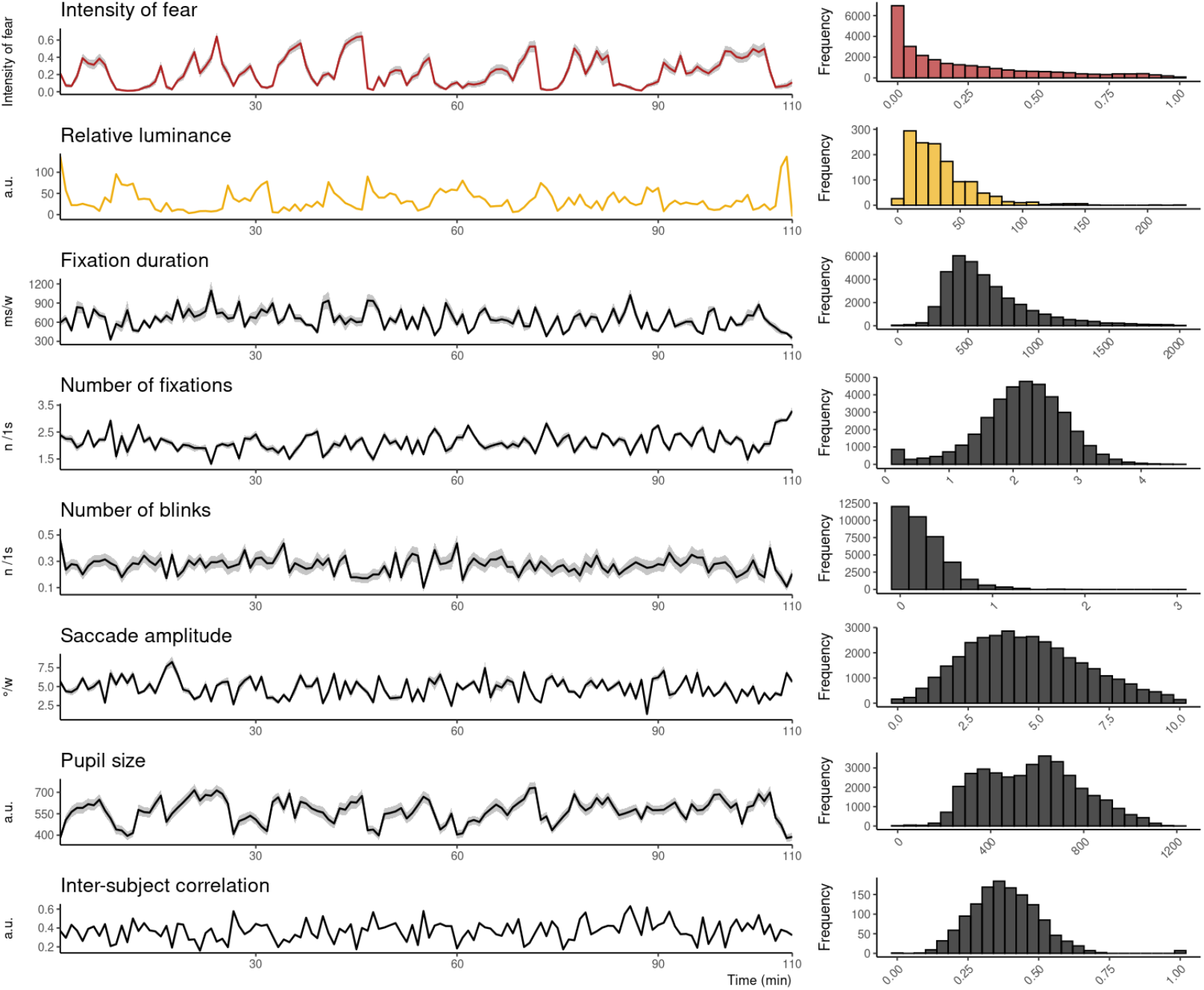
Mean (SEM) time series and distributions (over the whole time series) for intensity of fear, relative luminance and eye movement parameters during the movie in 5s time windows, and frequency distributions of the corresponding parameters. Inter-subject correlation measures the gaze synchronization among the subjects. Note: histograms show the data at subject level, which accounts for time windows with zero fixations and blinks.

Correlation analysis of timeseries revealed that fear ratings were positively associated with pupil size. Weak negative associations were found for blink and fixation rates and saccade amplitude, and correlations with e-ISC and fixation duration were weakly positive. Eye movement parameters showed the expected correlations with each other, with long fixation durations being associated with lower blink rates, number of fixations and saccade amplitudes, whereas e-ISC was positively associated with fixation duration and negatively associated with fixation and blinking rates and saccade amplitudes.

Notably, pupil size was positively associated with fear intensity, which in turn was negatively associated with relative luminance. This indicates that the associations between fear and pupil size (and possibly also other eye movement parameters) might be confounded with luminosity (e.g. scary scenes would consistently be darker). To address this, we regressed out the effect of luminance from each eye movement parameter and ran subject-wise regression models for fear and each eye movement parameter separately. Statistical testing of the resulting standardized betas (Figure 4) revealed that when the effect of luminosity on the eye movement time series was controlled for, fear was positively associated with larger pupil size (p = 2.9819*10^-9^) as well as negatively with shorter saccade amplitudes (p = 0.0025) and number of blinks (p = 0.0133). Associations between fear and fixation duration (p = 0.3059) and number of fixations (0.0738) were numerically positive but statistically nonsignificant.

**Figure 4.**
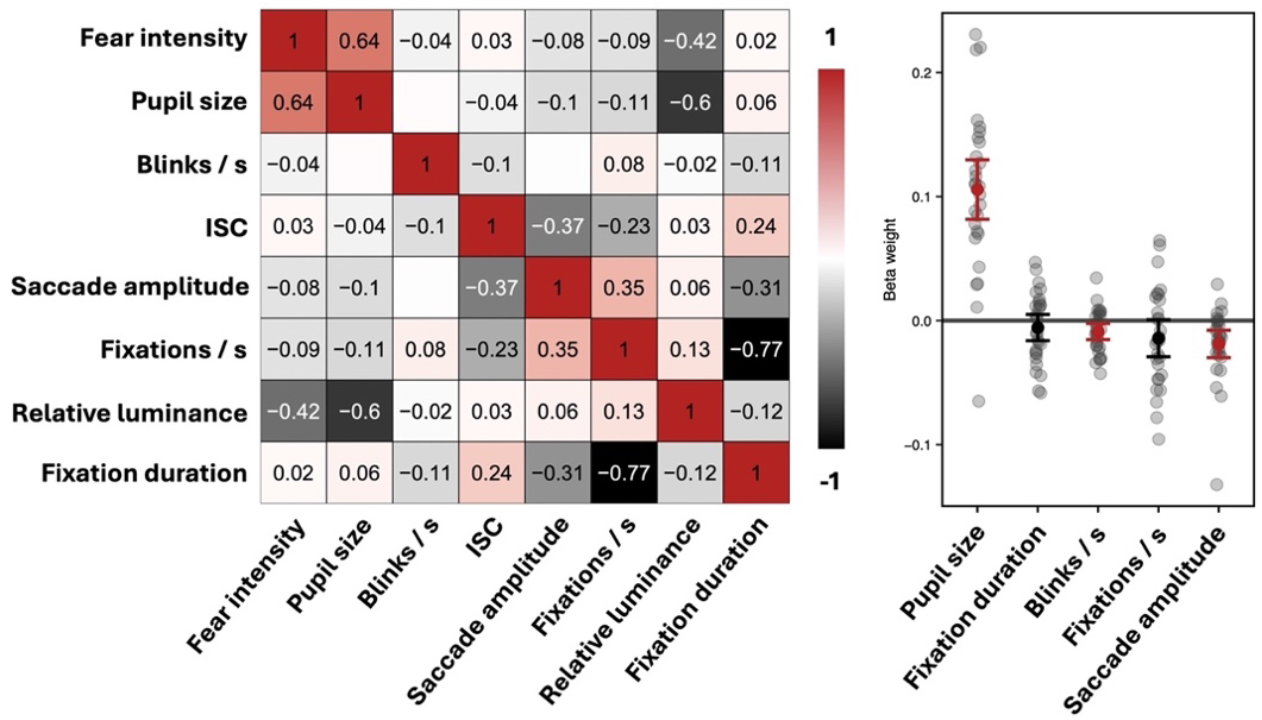
Left: Temporal associations (Pearson correlations) between eye movement parameters, fear and relative luminance across the whole movie. Only statistically significant associations (p < 0.05) are shown. Right: Subject-wise betas and their 95% CIs for the models between eye movement parameters and fear where the effect of luminance has been regressed out. Statistically significant (p < 0.05) effects are highlighted in red.

### Evoked eye movement responses to acute fear

Figure 5 shows the jump-scare evoked responses for the eye movement parameters. The highlighted area shows statistically significant time frames as found with cluster-based testing frameworks using 1000 permutations. Fixation duration and blink rate significantly decreased after the jump-scares whereas the number of fixations, saccade amplitude, and pupil size significantly increased. Inter-subject correlation did not show a significant change from baseline.

**Figure 5.**
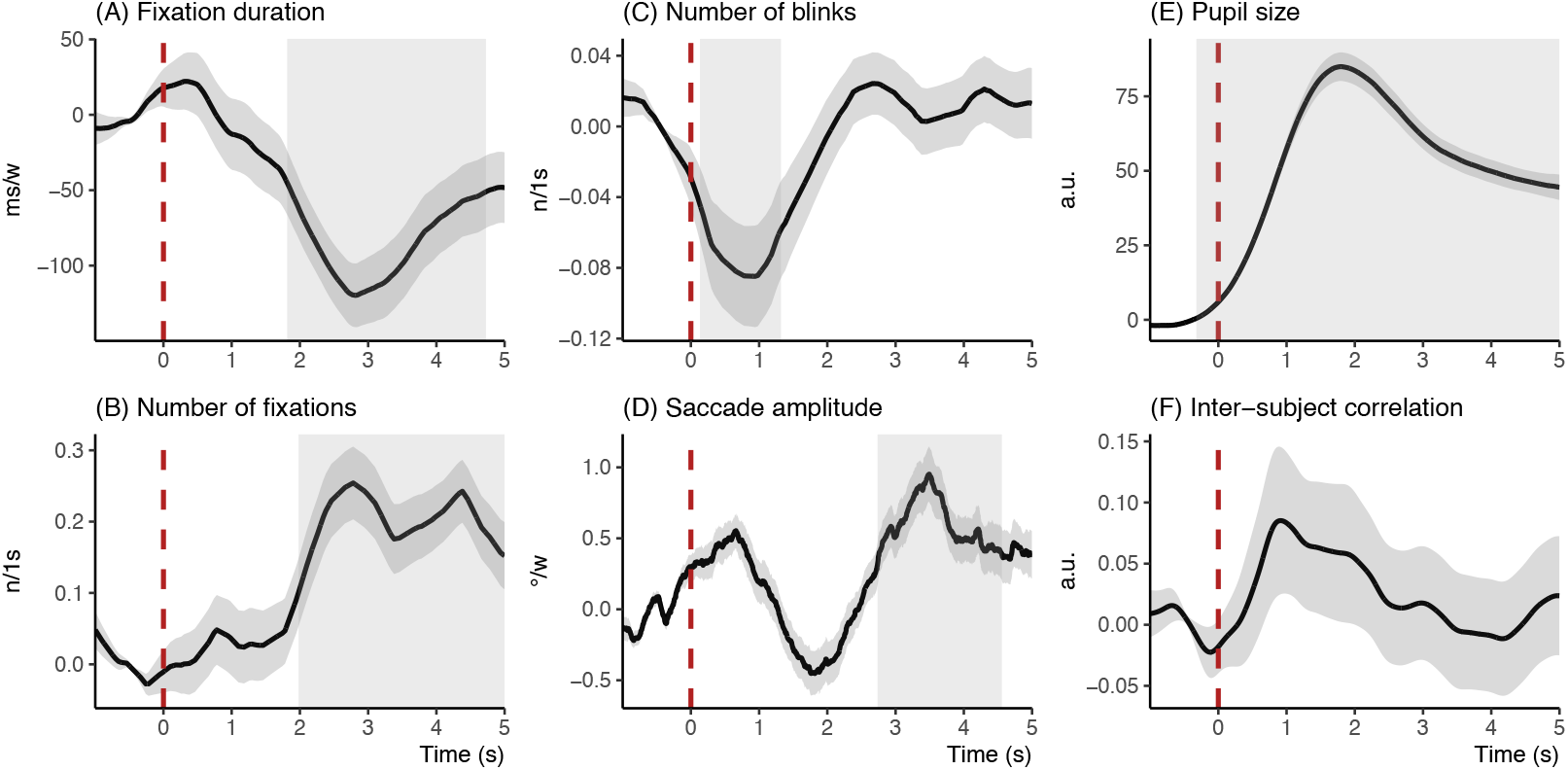
Normalized jump-scare-evoked responses (±SE, excluding inter-subject correlation) for eye movement parameters. The highlighted areas show the statistically significant timeframes (p < 0.05, two-tailed t-test). The red dashed line indicates the onset (0 ms) of the jump scares.

## Discussion

Our main finding was that both acute and sustained fear significantly modulate both visual sampling of dynamic scenes and affective arousal as indexed by pupil size. The analysis of the evoked responses triggered by the jump scares revealed that acute threats led to initial suppression of blinking and thus increased allocation of visual attention. This was followed by an increase in the spatial scanning of the scene, indicated by a decrease in the length of fixation, shortening of fixations and lengthening of saccades. Sustained, slower-frequency fluctuations in fear were in turn associated with shorter saccades and larger number of blinks, while both acute and sustained fear were associated with increases in pupil size, indicative of affective arousal. Altogether, our results show that both acute and sustained fear cause rapid reconfiguration and state-dependent changes in visual attention, supporting vigilance and effective scanning of the potentially threatening event.

### Evoked eye movement and pupillary responses to acute fear

Self-reports indicated that the horror movie evoked high and time-variable level of fear and pleasure, moderate level of disgust and tranquillity and low level of anger, confusion, joy and sadness (Figure 2). Although surprising, the simultaneously reported high levels of pleasure and fear are well in line with what we know from studies on “recreational fear”, where the thrills of horror movies and haunted houses arises from the fluctuation of the build-up and suspense of fear (Andersen 2020). Moreover, fear and pleasure share brain systems whose signals can be mixed under physically safe circumstances to promote experiences such as enjoyable fear (Nummenmaa 2024).

Acute fear events (jump scares) triggered a rapid increase in pupil size similar to the pupillary responses observed in the classical fear conditioning paradigms (Kluge, Bauer et al. 2011, Wiemer, Mühlberger et al. 2014, Korn, Staib et al. 2017). Accordingly, the rapid pupil dilation likely reflects an automatic startle response and heightened arousal. This arousal response was also accompanied with increased vigilance towards the scene. This was indicated by initial cessation in blinking immediately following the threat onset, which was followed by a short state of hypervigilance towards the scene, resulting in increased number of fixations, shortening of fixations and an increase in saccade lengths. This accords with studies showing that fear conditioning results in increased attention towards the conditioned stimuli (Klein, Ginat-Frolich et al. 2021, Rodriguez-Sobstel, Wake et al. 2023). However, our study revealed that attention capture and increased vigilance to threat onsets occurs also for unconditioned fear and reveals different time courses for the distinct components of the attentional responses. Blink rates, saccade lengths and fixation durations recovered to baseline levels during the analysed 5 s time window, whereas the effects of increased pupil size and fixation count extended beyond the analysis window. This indicates a biphasic change in attentional engagement. First, a rapid orienting response resulting in blinking cessation and initial “hypervigilance” consisting of short fixations accompanied with long saccades. Second, sustained changes in increased fixation count and arousal-dependent increases in pupil size.

Although prior work has found that attention is automatically captured by emotional stimuli across subjects (Nummenmaa, Hyona et al. 2006, Nummenmaa, Hyönä et al. 2009), the e-ISC analysis did not show any significant change in response to jump scares. In other words, participants’ eye movements did not become more similar during the acute threat episode. Although eye movements are more likely to orient towards threat than safety (Mulckhuyse, Crombez et al. 2013), the jump-scare events present in the movies do not necessarily contain a single, salient threat signal that would compete for attentional resources with the neutral or positive contents, as is common in eye-tracking paradigms where one emotional and one neutral scene simultaneously compete for attention. Although, we saw a numeric increase in e-ISC (about 0.1 units) during threat, it is likely that subjects were focusing on different areas of the screen during the acute threat, possibly trying to scan the scene to evaluate the current level and location of the threat (Bhushan and Asai 2018). It is also possible that the subjects intentionally avoided looking at the scariest content since 14/28 subjects reported frightening scene avoidance (see SI table1). Alternatively, it is possible that the slow-frequency e-ISC (based on spatial correlations computed over a time window) is simply not a fast enough metric for capturing the rapidly occurring fluctuations in eye movements. This explanation is however unlikely to be correct, as we did not observe any associations between e-ISC and sustained fear level (see below).

### Eye movement and pupillary responses to sustained fear

The analysis of the mean time series of the eye movement and pupil parameters revealed that pupil size was positively associated with fear but also with relative luminance. This accords with prior work indicating that pupillary responses are driven both by background luminosity and subjective emotion and arousal levels (Lowenstein, Feinberg et al. 1963, Bradley, Miccoli et al. 2008, Van Steenbergen, Band et al. 2011), and accords also with previously reported associations between acute threats and increased pupil size (Leuchs, Schneider et al. 2017). However, we also found a negative association between stimulus luminance and the intensity of self-reported fear. Darkness and the resulting uncertainty is a common method for generating suspense in horror movies as it sharpens the viewer’s senses, heightens alertness and primes expectations towards frightening events – perhaps because in our ancestral environment, darkness signalled increased vulnerability to predators or other unseen dangers (Nummenmaa 2024). Importantly, the association between fear level and pupil size remained significant even when we controlled for the effect of luminance. This confirms that this relationship was not merely due to luminosity differences between fearful and neutral scenes. Similar to acute fear, sustained fear was also associated with decreased blinking rate. Because blinking temporarily terminates visual input, studies have shown that decreased blinking is indicative of enhanced attentional engagement on the scene (Ranti, Jones et al. 2020). It is also possible that increased fear increases anticipatory protection of the eyes. Our data thus confirm that fear leads to this kind of engagement both at short and long time scales. However, unlike acute fear, sustained fear did not influence fixation rates or fixation durations, suggesting that the fear-dependent vigilant scanning is only induced by immediate rather than slow-frequency, “looming” threats.

Consistent with previous studies, we found that e-ISC remained moderately high throughout the experiment, indicating that a substantial proportion of the gaze patterns was stimulus-driven and thus occurring at the same spatiotemporal scales across observers (Henderson 2003, Lahnakoski, Glerean et al. 2014, Nummenmaa, Smirnov et al. 2014). Yet, similarly to acute fear, we only observed a weak correlation (r = 0.03) between e-ISC and experienced fear. All in all, these data suggest that despite the well-known attentional capture by emotional scene content, these transient changes in attention allocation do not translate to emotion-dependent synchronization of eye movements across longer time scales.

## Conclusion

We conclude that both acute and sustained fear evoked with Conjuring 2 horror movie modulate visual attention and arousal, as reflected in suppression of blinking and increase in pupil size. Abrupt threats additionally enhance attentional vigilance and detailed scanning of the environment, as indexed by shorter and more numerous fixations and longer saccades. Acute fear thus causes rapid reconfiguration and state-dependent changes in visual attention that support effective scanning of potentially threatening events, whereas sustained fear leads to increased arousal and vigilance, preparing the individual to respond to the potential threats in the environment.

## Acknowledgements

This study was supported by the Academy of Finland (grants #294897 and #332225), Sigrid Juselius Foundation, Turku University Foundation #080541, Instrumentarium Science Foundation (grants #200061 and #220030), Yrjö Jahnsson foundation #20217451, The Finnish Brain Foundation #20210063, The Päivikki and Sakari Sohlberg Foundation, The Paulo Foundation, and State research funding for expert responsibility area (ERVA) of the Turku University Hospital.

