## Supplementary information for "Attentional reconfiguration during acute and sustained fear"

### Content

1. **Questionnaires**
   1.1. Self-report questionnaires
   1.2. In-house questionnaire about attitude towards feeling of fear
2. **Correlation analysis**
   2.1. Correlations between mean eye movement parameters and sustained fear and anxiety, depression, emotion regulation, and empathy
   2.2. Correlations between jump-scare evoked eye movements and fear, anxiety, depression, emotion regulation, and empathy
   2.3. Correlations with eye-movement parameters and questionnaire scores
3. **Results**
   3.1. In-house self-reports
   3.2. Correlations between mean eye movement parameters and sustained fear and anxiety, depression, emotion regulation, and empathy
   3.3. Correlations between jump-scare evoked eye movements and fear, anxiety, depression, emotion regulation, and empathy
   3.4. Correlations with eye-movements and all the questionnaires
   3.5. Self-reports for horror movies
4. **Tables**
5. **Complete list of in-house questionnaires**
6. **References**

### 1. Questionnaires

#### 1.1 Self-report questionnaires

After watching the horror movie Conjuring 2, the subjects filled the Disgust Scale (DS-R)(Haidt, McCauley et al. 1994), modified by (Olatunji, Williams et al. 2007), Beck Depression Inventory (BDI-II) (Beck, Steer et al. 1988), Depression Anxiety Stress Scales (DASS) (Lovibond and Lovibond 1995), Emotion Regulation Questionnaire (ERQ) (Gross and John 2003), Interpersonal Reactivity Index (IRI) with four subcategories: Perspective Taking, Fantasy, Empathic Concern, and Personal Distress (Davis 1980), Short Psychopathy Rating Scale (SPRS), and The State-Trait Anxiety Inventory (STAI) (Spielberger CD 1970). These questionnaires were filled on the Redcap (Research Electronic Data Capture) platform (Harris, Taylor et al. 2009, Harris, Taylor et al. 2019).

#### 1.2 In-house questionnaire about attitude towards feeling of fear

To investigate how subjects were experiencing the feeling of fear, we developed in-house questionnaire to map 1) attitude towards the experience of watching horror movie / immersion, 2) fear regulation during the movie, and 3) physical symptoms. The questionnaire items are listed at the end of the SI file.

### 2. Correlation analysis

#### 2.1 Correlations between mean eye movement parameters and sustained fear and anxiety, depression, emotion regulation, and empathy

To test whether individual differences in the eye movement parameters are associated with emotional experience, emotion regulation, and empathy, pairwise correlations were computed between questionnaire scores and means of the eye movement time series like in main article.

#### 2.2 Correlations between jump-scare evoked eye movements and fear, anxiety, depression, emotion regulation, and empathy

To assess whether individual differences in personality scores were associated with evoked eye movement responses following jump scares, we first extracted the peaks by applying 6th degree polynomical curve with Matlab Curve Fitting toolbox to jump scare responses. Peaks and their latencies were then determined using function findpeaks. Finally, we correlated maximum peak values (minimum for saccade amplitude and fixation duration) with subjects' cognitive and emotion trait scores.

#### 2.3 Correlations with eye-movement parameters and questionnaire scores

We finally computed correlations between all the eye-tracking parameters, anxiety, depression, emotion regulation, empathy, physical symptoms, and in-house questionnaires. The correlation matrix was arranged by applying hierarchical clustering with R Studio ConsensusClusterPlus package (Wilkerson and Hayes 2010) that test with subset of data (80%) the probability if the variables belong the same cluster for 1000 times and used the consensus matrix as distance matric for hierarchical clustering. The optimal number of clusters by silhouette analysis was found 6 in Cluster package of R Studio. Finally, we combined the correlation and consensus matrices into one and marked the statistically ($p < 0.05$, Pearson) significant correlations with x.

### 3. Results

#### 3.1 In-house self-reports

The frequency distributions for the questionnaire items are shown in Figure S-1. Subjects experienced strong feelings of both horror and fear similarly. Majority of the subjects reported emphasizing with the characters' emotions and the movie, and they also reported high levels of immersions. The movie evoked also physical symptoms in the subjects. Most common symptoms were rapid heartbeat and dry mouth, indicative of autonomic activity.

The subjects reported using various emotion regulation strategies while viewing the movie. The most common means were focusing on neutral aspects, thinking about neutral things or simply just not trying to watch the movie. More specific, 7 subjects were thinking other things or reminded themselves that the movie is only a movie, 2 were anticipating, 1 added humor, 1 covered ears and 1 was playing with their fingers (Question 15).

Finally, majority of the subjects reported that horror movies make them feel excited, anxious, afraid, uncomfortable, nervous and tense (table S2).

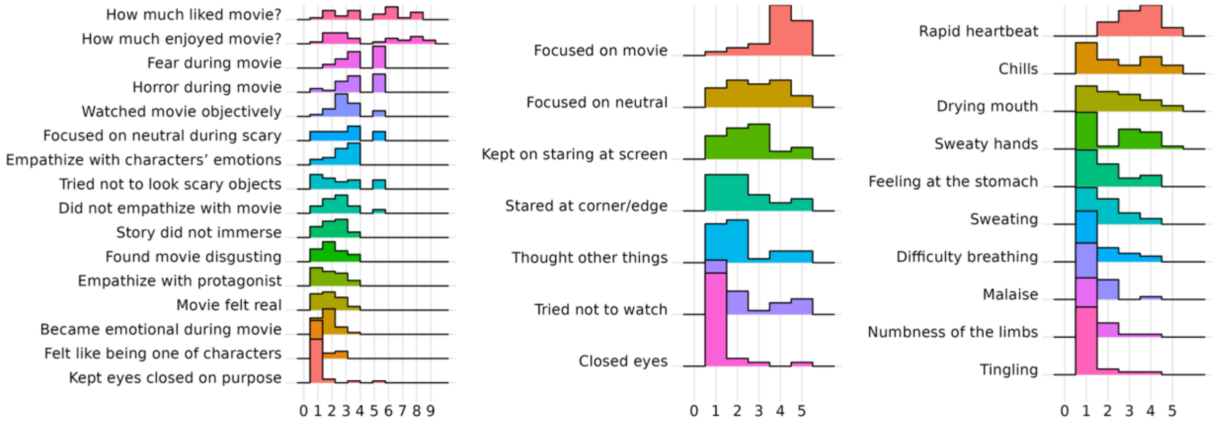


*Figure S-1: Subjective feelings, regulating the feeling of fear and physical symptoms while watching horror movie Conjuring 2. On the left column, questions "How much liked the movie?", and "How much enjoyed movie?" were asked on scale 1-10, the rest of the questions are on scale 1-5, where 1 = strongly disagree and 5 = strongly agree. Physical symptoms (middle) and fear regulations (right) were on scale 1-5, where 1 = not at all and 5 very much.*

### 3.2 Correlations between mean eye movement parameters and sustained fear and anxiety, depression, emotion regulation, and empathy

Associations between means of the eye movement parameters and questionnaire scores are in Figure S-2. In general, questionnaire scores were not strongly associated with eye tracking parameters. Questionnaires measuring stress, anxiety or depression suggested positive association with fixation duration and negative association with blink and fixation rates, but the correlations were not statistically significant. However, blink rate had statistically significant and negative association with anxiety (STAI) and secondary psychopathy score. High IRI fantasy scores were positively associated with fixation duration and negative association with fixation rate.

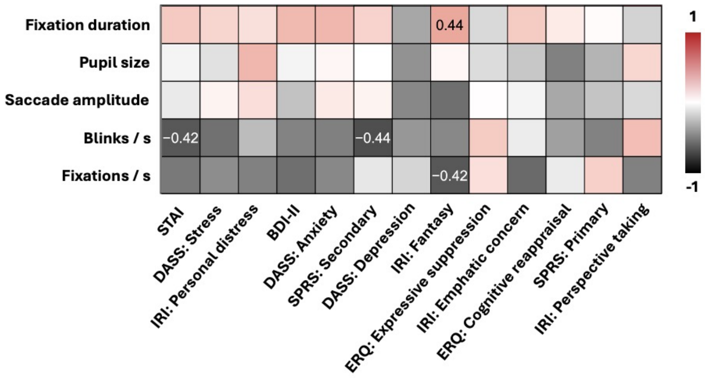


*Figure S-2: Associations (Pearson correlation) between mean eye movement parameters and questionnaire scores. Only statistically significant associations (p < 0.05) are shown. BDI-II = Beck Depression Inventory, DASS = Depression Anxiety Stress Scales,*

#### 3.3 Correlations between jump-scare evoked eye movements and fear, anxiety, depression, emotion regulation, and empathy

The correlation matrix is shown in Figure S-3. Some of the emotional and cognitive traits had significant correlations with the latency or peak height (minimum values for saccade amplitude and fixation duration) of the eye tracking responses to jump scare. Questionnaires measuring anxiety, depression or stress had negative association with peak latency in fixation duration (the more anxious the patient is the longer the fixation duration was) and positive association with the peak latency in fixation rate (the more anxious the patient was the less they took fixations). Maximum amplitude of fixation rate had positive association with primary psychopathy scores and negative association with personal distress; the higher the score in primary psychopathy the more fixations and the more empathy the less fixations. Emphatic concern had negative association with minimum amplitude of fixation duration and Perspective taking positive association with maximum amplitude of pupil size. In terms of emotion regulation, none of the correlations were statistically significant.

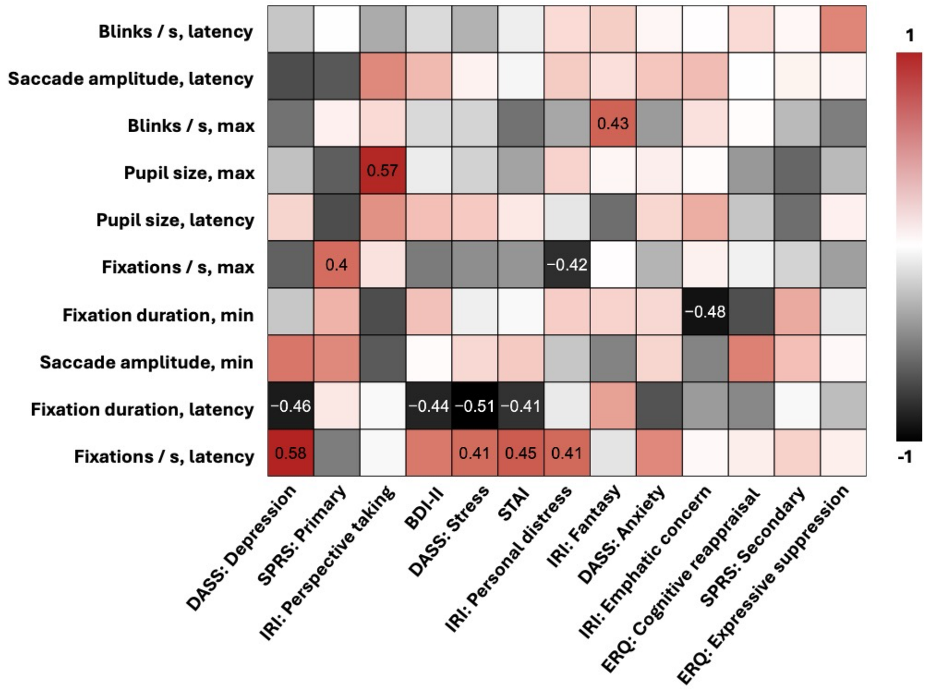


*Figure S-3: Associations (Pearson correlations) between the personality scores and peak height and latency for the post-jump-scare evoked responses. Only statistically significant associations ($p < 0.05$) are shown.*

#### 3.4 Correlations with eye-movements and all the questionnaires

The correlation matrix (Figure S-4) revealed an association between watching and liking horror movies. The more the subject enjoyed the movie, the more they liked the movie and the more they reported liking watching horror. The less the subject enjoyed the movie the more they tried to avoid watching it, to think other things, to stare at corner or edges or to focus on neutral objects.

Physiological symptoms were associated with immersion, horror and avoidance of watching. Enjoying the movie had a negative association with self-reported rapid heartbeat or amount of fear during the movie. Feelings at the stomach had a positive association with finding the movie disgusting. Feelings at the stomach correlated significantly keeping eyes closed while viewing the film, staring at the corners or edges, focusing on neutral issues, and trying not to watch. Regarding emotion regulation, avoiding watching the movie had positive association with thinking other things. Thinking other things was positively associated with self-reported heartbeat, fear or horror during the movie, higher emphatic concern or higher points in IRI fantasy. Finally, the more the subject immersed themselves in the movie, the more horror they felt and the more they tried to ease the feeling.

#### 3.5 Self-reports for horror movies

Self-reported preferences for horror movies are shown in tables S3-S6.

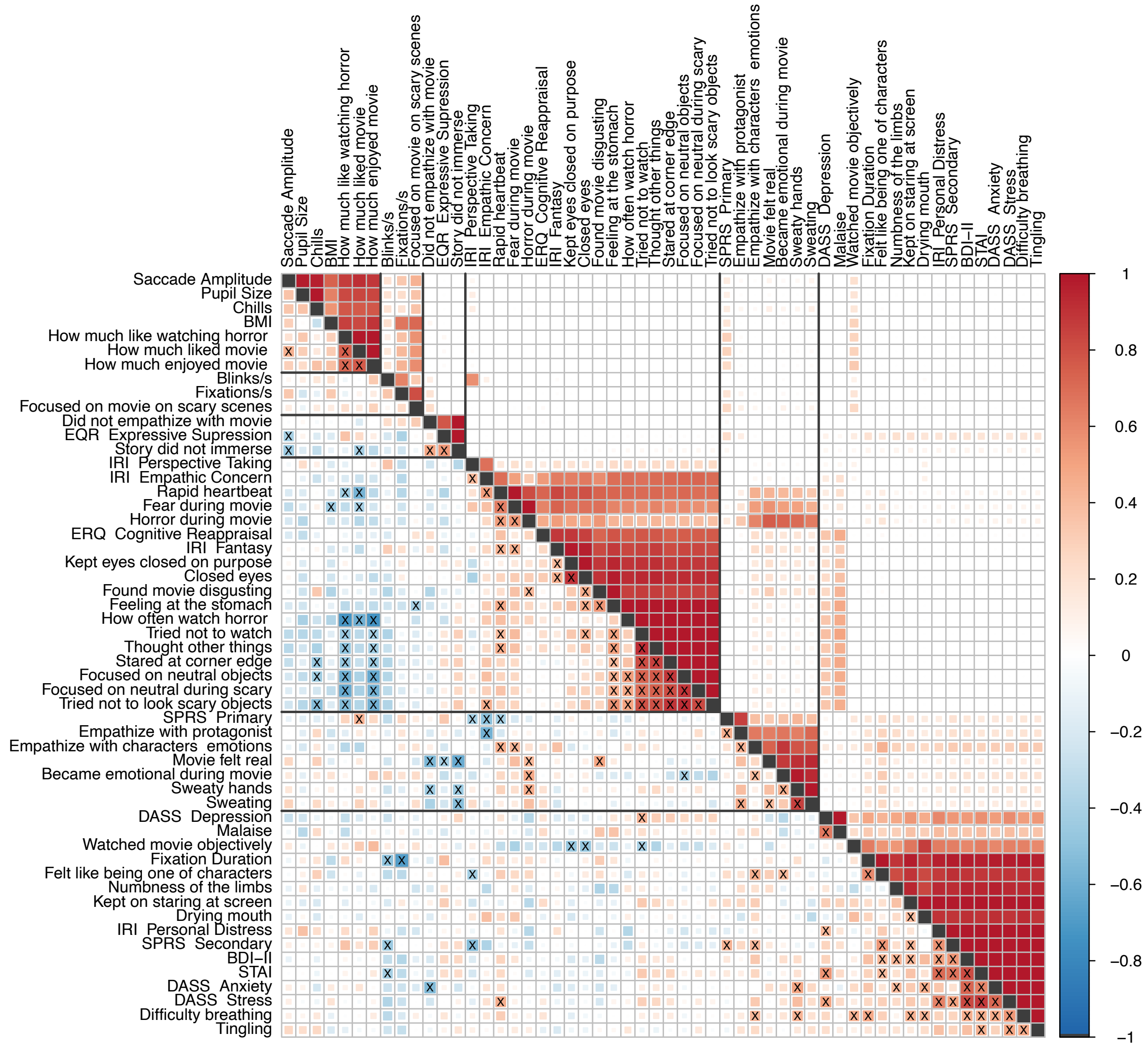


*Figure S-4: UPPER TRIANGLE: consensus matrix with hierarchical clustering for organizing the correlation matrix. LOWER TRIANGLE: Associations (Pearson correlations) of mean eye-tracking parameters, personality traits, and experiences of watching horror movie Conjuring. X marks statistically significant correlations (p <0.05). BDI-II = Beck Depression Inventory, DASS = Depression Anxiety Stress Scales, ERQ = Emotion Regulation Questionnaire, IRI = Interpersonal Reactivity, SPRS = Short Psychopathy Rating Scale, STAI = The State-Trait Anxiety Inventory. NB: the scale is only 0-1 for consensus matrix.*

*Table 1: 14/28 subjects reported to stare the corner, center, edge or somewhere else in the screen during frightening scenes.*

| Frightening scenes avoidance | n |
|---|---|
| Corner | 4 |
| Center | 0 |
| Edge | 3 |
| Corner and edge | 4 |
| Somewhere else | 3 |

*Table 3: Table shows how often 28 participants were watching movies and horror movies.*

| | Frequency of watching | |
|---|---|---|
| | movies | horror movies |
| Daily | 0 | 0 |
| Once a week | 0 | 0 |
| Once in 2 weeks | 11 | 0 |
| Once a month | 7 | 3 |
| Once in 3 months | 7 | 5 |
| Rarely | 3 | 20 |

*Table 2: The 28 participants reported all the feelings horror movies made them feel.*

| Horror film induced feelings | n |
|---|---|
| Excited | 26 |
| Anxious | 26 |
| Afraid | 25 |
| Uncomfortable | 25 |
| Nervous | 21 |
| Tense | 19 |
| Frustrated | 17 |
| Disgusted | 14 |
| Surprised | 13 |
| Puzzled | 12 |
| Paranoid | 11 |
| Curious | 11 |
| Confused | 10 |
| Interested | 8 |
| Cheerful | 7 |
| Insecure | 6 |
| Dull | 6 |
| Relieved | 5 |
| Pleasure | 4 |
| Joyful | 3 |
| Courageous | 3 |
| Numb | 2 |
| Nauseous | 2 |
| Escapist | 1 |
| Derisive | 1 |

*Table 4: 28 subjects reported what kind of movies they usually watch the most (1st), the second to most (2nd) and third to most (3rd), and their favourite movie genre.*

| | **Usually watched movies** | | | **Favourite genre** |
|---|---|---|---|---|
| | 1st | 2nd | 3rd | 1st |
| Drama | 11 | 1 | 2 | 10 |
| Documentary | 4 | 1 | 5 | 4 |
| Comedy | 2 | 7 | 6 | 4 |
| Thriller | 2 | 3 | 2 | 3 |
| Adventure | 2 | 2 | 2 | 3 |
| Romantic | 2 | 2 | 2 | 1 |
| Horror | 1 | 2 | 3 | 1 |
| Action | 1 | 6 | 1 | 1 |
| Sci-fi | 0 | 1 | 1 | 0 |
| Musicals | 0 | 0 | 1 | 1 |
| Na | 3 | 3 | 3 | - |

*Table 5: Table presents how much the participants like to watch movies or like horror movies.*

| | **watching movies** | **horror movies** |
|---|---|---|
| **Liking*** | n | n |
| 1 | 0 | 4 |
| 2 | 1 | 5 |
| 3 | 6 | 4 |
| 4 | 11 | 1 |
| 5 | 8 | 3 |
| 6 | - | 1 |
| 7 | - | 4 |
| 8 | - | 3 |
| 9 | - | 3 |
| 10 | | 0 |
| NA | 2 | - |

*(1= not at all, 5 or 10 = very much)

*Table 6: 28 participants reported all the types of horror movies they found scary.*

| **Scariest horror films** | **n** |
|---|---|
| Psychological horror | 27 |
| True crime | 15 |
| Torturing | 14 |
| Survival horror | 12 |
| Supernatural | 10 |
| Crime horror | 9 |
| Blood and bloodshed | 6 |
| Teen horror | 1 |
| Monsters | 1 |
| Gothic horror | 1 |
| Sci-fi | 0 |

**Survey: Conjuring 2**

Assess the following claims based on the film. Use the following scale:
(1 = Strongly disagree 5 = Strongly agree)

1. I was experiencing fear while watching the movie 1 2 3 4 5
2. I was experiencing horror while watching a movie 1 2 3 4 5
3. I found the movie disgusting 1 2 3 4 5
4. I watched the movie objectively 1 2 3 4 5
5. I kept my eyes closed on purpose 1 2 3 4 5
6. I tried to focus on neutral issues during the scary scenes 1 2 3 4 5
7. The movie felt real 1 2 3 4 5
8. I could easily empathize with the protagonist 1 2 3 4 5
9. I tried not to look at the scary objects on the screen so that I would not be afraid 1 2 3 4 5
10. I did not empathize strongly with the movie 1 2 3 4 5
11. I became emotional during the movie 1 2 3 4 5
12. I felt like I was one of the characters in the story 1 2 3 4 5
13. I empathized with the characters' emotions 1 2 3 4 5
14. The story of the movie did not immerse me 1 2 3 4 5
15. When I was really scared during the movie, I tried to ease the feeling?

    No Yes

    How?______________________________________________________

16. On the scale of 1 -10 (1 = Not at all, 10 = Very much), how much did you <u>like</u> the movie?

    1 2 3 4 5 6 7 8 9 10

17. On the scale of 1 - 10 (1 = Not at all, 10 = Very much), how much did you <u>enjoy</u> the <u>experience</u> of watching the movie? 1 2 3 4 5 6 7 8 9 10

18. Did you do any the following things while watching the movie?

    1= Not at all, 5 = Very much

1. I focused on neutral issues on the screen during frightening scenes 1 2 3 4 5
2. During the frightening scenes, I stared at the **corner/center/edge** of the screen, or somewhere else (underline where) 1 2 3 4 5
3. I thought about other things during the scary scenes 1 2 3 4 5
4. I kept on staring at the screen when I stopped focusing on the movie 1 2 3 4 5
5. I focused on the movie during the scary scenes 1 2 3 4 5
6. I tried not to watch the scary scenes 1 2 3 4 5
7. I closed my eyes during the scary scenes 1 2 3 4 5

19. Did you experience following physical sensations during the movie? (1 = Not at all, 5 = Very much)

1. Sweaty hands 1 2 3 4 5
2. Feeling at the bottom of the stomach 1 2 3 4 5
3. Sweating 1 2 3 4 5
4. Malaise 1 2 3 4 5
5. Chills 1 2 3 4 5
6. Rapid heartbeat 1 2 3 4 5
7. Drying mouth 1 2 3 4 5
8. Difficulty breathing 1 2 3 4 5
9. Numbness of the limbs 1 2 3 4 5
10. Tingling 1 2 3 4 5

The following questions are about your general opinions regarding films. Estimate the validity of the following claims.

20. On the scale of 1 - 5 (1= Not at all, 5 = Very much),
    How much do you like movies? 1 2 3 4 5

21. How often do you watch the movies?

| | | | | | |
|---|---|---|---|---|---|
| Daily | Once a week | Once in two weeks | Once a month | Once in three months | Rarely |

22. What kind of movies you usually watch? Place the numbers 1, 2 and 3 for the movie genre you watch the most (1= the most):

| | | |
|---|---|---|
| Documentary | Horror | Thriller |
| Drama | Romantic | Western |
| Comedy | Adventure | Sci-fi |
| Musicals | Action | |

23. What is your favorite movie genre?

| | | |
|---|---|---|
| Documentary | Horror | Thriller |
| Drama | Romantic | Western |
| Comedy | Adventure | Sci-fi |
| Musicals | Action | |

24. Do you like horror films? No Sometimes Yes

25. How much do you like watching horror films? (1= Not at all, 10 = Very much)

1 2 3 4 5 6 7 8 9 10

26. How often do you watch horror films?

| | | | | | |
|---|---|---|---|---|---|
| Daily | Once a week | Once in two weeks | Once a month | Once in three months | Rarely |

27. How do horror films make you feel? (select all suitable)

| | | | |
|---|---|---|---|
| Tense | Confused | Puzzled | Paranoid |
| Excited | Nauseous | Courageous | Derisive |
| Afraid | Frustrated | Curious | Surprised |
| Anxious | Relieved | Escapist | Uncomfortable |
| Nervous | Dull | Interested | |
| Disgusted | Cheerful | Joyful | |
| Pleasurable | Insecure | Numb | |

28. What types of the horror films are the scariest? (select all suitable)

| | | |
|---|---|---|
| Psychological horror | Torturing | Monsters |
| Blood and bloodshed (gore) | True crime | Sci-fi |
| Survival horror | Supernatural | Gothic horror |
| Crime horror | Teen horror | |

Questions about the plot:

1. What are Loraine's and Ed Warren's job descriptions?
   What kind of methods do they use at their work?
2. How and when does Janet become possessed?
3. How does the story of the family conclude?
